# Cardiac Extracellular Matrix derived from younger animals stabilizes macrophage inflammatory polarization through modulation of the interferon signaling pathway

**DOI:** 10.64898/2026.09.08.750183

**Authors:** Keely Nugnes, Grace Costello, Nora McCarey, Ella Canas, Lauren D Black

## Abstract

Neonatal mammals exhibit a remarkable capacity for cardiac repair that declines rapidly after birth and continues to diminish with age. This loss of regenerative potential is accompanied by age-dependent changes in macrophage behavior, shifting from pro-regenerative to pro-fibrotic responses following injury. We investigated whether extracellular matrix (ECM)-derived matrikines from different developmental ages regulate macrophage polarization. Cardiac ECM was isolated from neonatal (P2), adolescent (6-week), and adult (16-20-week) rat hearts through decellularization and pepsin digestion. ECM peptides were adsorbed onto tissue culture surfaces, and murine RAW 264.7 macrophages were cultured on ECM-coated or uncoated controls. After 24 hours, macrophage phenotype and gene expression were assessed using marker analysis and RNA sequencing. Macrophages cultured on neonatal cardiac ECM displayed reduced inflammatory activation and maintained expression of pro-reparative markers, even in the presence of inflammatory stimuli. Transcriptomic analysis revealed a distinct ECM-driven phenotype that differed from canonical M1 and M2 polarization states. Gene ontology analysis demonstrated significant downregulation of pathways associated with antiviral responses, immune activation, and interferon signaling in macrophages exposed to neonatal ECM. These effects were not observed with adolescent or adult ECM. Collectively, these findings suggest that neonatal ECM matrikines suppress interferon-mediated inflammatory signaling, potentially contributing to the reduced inflammation and enhanced repair observed following myocardial injury in neonatal hearts.

## INTRODUCTION

Macrophages, a type of phagocytic immune cell, play a primary role in regulating wound healing. Rather than existing as a single functional entity, macrophages adopt a range of phenotypic states in response to environmental cues, with these states critically shaping whether tissue healing proceeds toward regeneration or chronic inflammation and fibrosis. In the initial, inflammatory stage of wound healing, macrophages release inflammatory factors and phagocytose necrotic debris. Inflammation is resolved during the pro-reparative stage, in which macrophages stimulate extracellular matrix (ECM) production, cell proliferation, and angiogenesis ^1^. The transition from the initially pro-inflammatory phenotype to the pro-reparative phenotype is crucial to healthy wound healing, resolution of inflammation, and restoration of tissue homeostasis ^2^.

In experimental settings, macrophage phenotypes have often been described using the simplified framework of pro-inflammatory “M1” and pro-reparative “M2” polarization states. Classical “M1” polarization is induced by E. coli lipopolysaccharide (LPS) and interferon gamma (IFN-□) stimulation, whereas the “M2” like polarization is traditionally achieved through linterleukin-4 (IL-4) stimulation ^2^. While this classification had provided a useful foundation for studying macrophage biology, it is now well recognized that macrophage activation in vivo is far more heterogeneous and context dependent. Rather than discrete states, increasing evidence has shown M1 and M2 macrophages represent extremes along a continuous spectrum of activation, with intermediate and mixed phenotypes emerging in response to complex, dynamic tissue environments ^3^. Importantly, macrophage function is shaped not only by soluble cytokines, but also by local structural and biochemical cues, such as tissue stiffness ^4^, metabolic activity ^5^, and epigenetic regulation ^6^, suggesting that tissue-specific factors may play a key role in regulating macrophage polarization during repair.

The importance of macrophage phenotype in shaping tissue remodeling is particularly evident in the heart following myocardial infarction (MI). Adverse tissue remodeling following MI leads to progressive ventricular dysfunction and remains a major cause of chronic heart failure. In the adult mammalian heart, the limited proliferative capacity of cardiomyocytes results in the formation of a fibrotic scar that permanently compromises cardiac function ^7^. In contrast, neonatal mammals retain a transient capacity for cardiac regeneration, enabling restoration of myocardial structure without permanent scar formation ^8^. Macrophages play a critical role in this response, as removal of macrophages from the neonatal heart abolished regeneration and results in fibrotic repair, demonstrating that macrophages are required for regenerative healing ^9^. Conversely, in the adult heart, macrophages actively contribute to fibrosis and scar formation following injury ^10^. These observations highlight the context-dependent nature of macrophage behavior and raise the possibility that age-specific cues within the cardiac microenvironment instruct macrophages towards regenerative or fibrotic phenotypes.

One key component of the tissue microenvironment that undergoes dynamic remodeling during injury and repair is the extracellular matrix. Tissue remodeling during homeostasis and wound healing is initiated, in part, by ECM degradation mediated by proteases, released from cells such as macrophages and fibroblasts, break ECM proteins down into small peptides that have the ability to give unique signals to cells relating to tissue remodeling. The term “matrikines” has been commonly used to describe these peptides ^11^. Increasing evidence demonstrates that signals derived from these peptides can influence a range of cellular processes, including proliferation, migration, and differentiation ^12^. Importantly, these signals can also modulate immune cell behavior, including macrophage phenotype and polarization ^13^. Because ECM composition and structure change dramatically across developmental stages, ECM derived from tissues of different developmental ages may encode distinct biochemical information capable of differentially instructing macrophage responses.

The purpose of this study was to investigate how the developmental age of cardiac ECM influences macrophage phenotype and polarization. We hypothesized that neonatal cardiac ECM, which supports regenerative healing in vivo, would promote a pro-reparative, M2-like macrophage phenotype relative to adult ECM. Consistent with this hypothesis, macrophages cultured on neonatal ECM exhibited increased proliferation and attachment and adopted a distinct M2-like phenotype characterized by elevated expression of pro-reparative markers. Notably, this phenotype was maintained even in the presence of inflammatory stimuli, suggesting that neonatal ECM provides dominant immunomodulatory cues. Together, these findings demonstrate that developmental age-specific ECM signals can direct macrophage behavior and highlight the therapeutic potential of neonatal cardiac ECM as in immunomodulatory biomaterial for the treatment of cardiac injury and disease.

## METHODS

### Heart Harvest from Neonatal and Adult Rats

All animal procedures were performed in accordance with the Institutional Animal Care and Use Committee at Tufts University and the NIH Guide for Care and Use of Laboratory Animals. Female adult (∼16-20 weeks) and young adult (6 weeks) Sprague Dawley rats were euthanized using CO_2_ followed by the harvest of the heart. Neonatal rats pups (postnatal day 2) were euthanized by conscious decapitation, followed by heart harvest.

### Decellularization and Solubilization of Cardiac ECM

Hearts were decellularized and solubilized using protocols previously described in the literature (^12, 14^). Ventricular tissue was minced and decellularized using 0.1% or 1% (w/v) sodium dodecyl sulfate (SDS) in deionized water for neonatal and adult hearts, respectively, with gentle shaking for two to five days, until decellularization was complete and hearts were no longer opaque. Hearts were then washed with deionized water for 24 hours, then flushed three times with large amounts of deionized water to remove any remaining SDS. ECM was frozen at -20°C overnight, and then lyophilized for 24-48h, until completely dry.

ECM samples were minced and solubilized using pepsin at a 10:1 tissue to pepsin ratio in 0.1M HCl. Samples were mixed with a magnetic stir bar at ∼800rpm at room temperature (RT) until no solid tissue pieces were visible and solution was homogenous, approximately 3 hours for neonatal tissue and 1-2 days for adult tissue. 1M NaOH was added to bring the solution past pH 8, to permanently deactivate the pepsin, and then neutralized. ECM was brought to a final concentration of 5 mg ml^-1^, aliquoted, and stored at -20°C for future use, or used immediately.

### Preparation of ECM-Coated Substrates

Solubilized ECM was allowed to adsorb onto 24-well tissue culture polystyrene dishes at 100 ug cm^-2^ and allowed to air dry for 1-2 days in a sterile tissue culture cabinet. Excess solution was aspirated and wells were rinsed with sterile phosphate buffered saline (PBS) with 1% penicillin-streptomycin (P-S) immediately prior to use.

### RAW 264.7 mouse macrophage culture and polarization

RAW 264.7 macrophages were expanded in Dulbecco’s Modified Eagle Medium (DMEM) supplemented with 10% fetal bovine serum (FBS) and 1% P-S in a humidified incubator at 37°C and 5% CO_2_. Macrophages were plated in ECM treated 24-well plates at 50,000 cells/well in DMEM with 2% FBS and 1% P-S. Uncoated wells were used as M0 controls. Macrophages were polarized towards an M1 phenotype by adding 100 ng mL^-1^ LPS and 20 ng mL^-1^ IFN-γ or an M2 phenotype by adding 30 ng mL^-1^ IL-4 to the media. Macrophages were cultured on ECM-coated substrates or in polarization media for 24 hours before further analysis. To stimulate cells with interferon molecules, cells were incubated in either 20 ng mL^-1^ IFN-γ or 1,000 U mL^-1^ of interferon-β (IFN-β).

### Induced Pluripotent Stem Cell Differentiation

Human induced pluripotent stem cells (iPSCs) were maintained and differentiated using protocols commonly used in the literature ^15^. To differentiate iPSCs into cardiomyocytes, iPSCs were maintained on Matrigel coated plates in Roswell Park Memorial Institute Medium (RPMI) with B27 supplement minus insulin and 6μM CHIR99021. On day 2, cells were rested with RPMI-B27 minus insulin for 24 hours, then 2μM C59 was added for two days. On day 7, cells were switched to RPMI containing B27 supplement with insulin. Cells were maintained until day 16 with media changes every other day. On day 16, iPSC-CM were frozen for future experiments.

To differentiate iPSCs into cardiac fibroblasts (iPSC-CF), iPSCs were maintained on Matrigel coated plates in RPMI supplemented with B27 media without insulin and 6μM CHIR9902. On day 3, media was replaced with RPMI supplemented with B27 supplement without insulin and 2μM C59. On day 6, cardiac progenitor cells were re-plated at 20,000 cells/cm^2^ in Advanced DMEM with GlutaMAX media containing 5μM CHIR, 2μM retinoic acid, 5μM ROCK inhibitor, and 1% FBS. After 24 hours, media was replaced without ROCK inhibitor. Media was changed every other day for 4 days. On day 11, cells were passaged at a 1:3 ratio and plated on Matrigel coated plates in Advanced DMEM with GlutaMAX supplemented with SB431541. Cells were maintained as epicardial cells, with media changes every other day for at least three days. To differentiate into iPSC-CF, cells were passaged and seeded at 10,000 cells/cm^2^ in Fibroblast Growth Medium supplemented with 20 ng mL^-1^ FGF2 and 10μM SB431541. Cells were maintained for 4 days with one media change, then re-plated at a 1:3 ratio. On day 20, iPSC-CF were considered fully differentiated. Cells were passaged a maximum of 4 times or used immediately for experiments.

### Cardiomyocyte Conditioned Media Studies

Macrophages were grown on ECM-coated substrates for 24 hours to develop macrophage conditioned media (MCM). Media was filtered in a 0.2μm filter, to remove any cells still in suspension, and used immediately.

iPSC-CM were thawed and seeded at a density of 200,000 cells/well in a 24 well-plate and maintained for 4 days, until iPSC-CM exhibited strong, spontaneous beating. On day 5, media was changed to serum-free RPMI for 3h, at which point media was replaced by 50:50 B27 media and MCM. To determine proliferation, iPSC-CM were incubated for 24 hours in conditioned media, then assessed for their expression of Ki-67. For cell survival, iPSC-CM were incubated in MCM for 1h, then media was spiked with 200μM H_2_O_2_ for 3 hours, at which point cells were fixed and underwent a TUNEL assay to determine cell death.

### Fibroblast Conditioned Media Studies

iPSC-CF were passaged at least once before seeding at 10,000 cells/well in a 24 well plate in Fibroblast Growth Media (FGM). After 24h in culture, iPSC-CF were serum starved for 24h, then stimulated with 50:50 serum free media and MCM, with or without the addition of 10 ng mL^-1^ TGF-β. After 48h, cells were fixed and stained for their expression of α-Smooth muscle actin (α-SMA) and Ki-67.

### Fluorescent Staining and Image Analysis

Cells were fixed with 4% paraformaldehyde at 4°C for 10 minutes. For stains targeting intracellular epitopes, samples were permeabilized with 1% Triton-X 100 for 10 minutes at RT. Samples were blocked with 1% bovine serum albumin (BSA) and 5% donkey serum in PBS overnight at 4°C, then stained overnight with anti-Ki-67 (Abcam, ab15580), anti-CD206 (Abcam, ab64693), anti-CD86 (Abcam, ab119857), anti-CD163 (Thermo Fisher, 16646-1-AP), anti-iNOS (Abcam, ab178945), anti-α-SMA (Abcam, ab7817), or anti-α-actinin (Sigma Aldrich, 7811) in 1% BSA in PBS at 4°C. Next, samples were incubated with either donkey anti-rabbit or donkey anti-rat secondary antibodies at 1:1000 dilution in 1% BSA with 1 mg mL^-1^ Hoechst 33342 for 2h. For morphological analyses, cells were also incubated with phalloidin 568 (Invitrogen) at 1:400 dilution.

Samples were imaged on a Keyence BZ-X710 series microscope and analyzed using custom Cell Profiler (Broad Institute) pipelines. To determine macrophage expression of phenotypic markers, the total number of cells and the number of cells expressing stained epitopes were counted in order to determine the number of Ki-67, CD86, CD206, CD163, or iNOS positive cells. For morphological analyses, cell shapes were identified using the cytoskeletal stain phalloidin, and measured for their perimeter (P), area (A), minor axis length, and major axis length. Cell circularity was calculated using 4π A P^-1^ and cell elongation calculated as the major axis to minor axis length ratio.

### Enzyme Linked Immunosorbent Assay

Macrophage cytokine release was measured after 24 hours of culture on ECM-coated substrates or in polarization media. The amount of secreted IL-6, IL-10, TGF-β, and IL-1β were measured using enzyme linked immunosorbent assay (ELISA), per manufacturers instructions (Invitrogen). In short, a capture antibody was allowed to adsorb onto high-protein binding plastic overnight. Media samples were added in triplicate and target cytokines were allowed to attach to antibodies overnight. Then, a biotinylated detection antibody was added, followed by streptavidin-conjugated horseradish peroxidase. A colorimetric reaction with tetramethylbenzidine was allowed to carry out for 15 minutes before stopping with 1M HCl. A plate reader was used to measure absorbance at 450nm with 620nm for wavelength subtraction.

### RNA Preparation and Sequencing

Macrophages were grown on ECM-coated substrates or in polarization media for 24 hours, at which point cells were lysed and RNA was isolated using an RNeasy Mini Kit (Qiagen), per manufacturers instructions. RNA was quantified on a Nanodrop 2000 UV-Vis Spectrophotometer (Thermo Fisher Scientific) and stored at -80°C until further use.

Purified RNA samples were delivered to Tufts University Genomics Core Facility for quality certification, cDNA library construction (Illumina TruSeq Stranded mRNA), and sequencing. Sequencing was performed on an Illumina NovaSeq X Plus (100-bp single-end lane). Raw data was trimmed using Trimmomatic ^16^ and quality validated using FASTQC ^17^. Data was then loaded into STAR (v. 2.7.11b) ^18^ to align sequences with the mouse genome GRCm38.p4 (from the Gene Code for the Encyclopedia of DNA Elements (GENCODE) database). STAR aligned reads were stored as BAM files.

### Differential Gene Expression Analysis

Read counts were generated and generalized fold change was generated from aligned sequencing data using GFOLD ^19^. For samples grown on ECM-coated substrates or in polarization media, the M0 group was assigned as the control condition for analysis. Output files were loaded into R Studio for further analysis. Principal component analysis was performed with the plotPCA function and normalized gene expression data from all groups was output into a hierarchial clustering heatmap with the pheatmap function in R. Principal Component Analysis (PCA) plots were generated using the prcomp and ggplot functions. Gene enrichment analysis was done using the BiocManager package in R, which includes functions for comparing top differentially expressed genes, as determined by genes with a fold change greater than 1 or less than -1, against Gene Ontology (GO) ^20^ and Kyoto Encyclopedia of Genes and Genomes (KEGG) ^21^ databases.

### Statistical Analysis

Statistical significance was determined using dimensionally appropriate analysis of variance (ANOVA) tests, followed by Tukey’s multiple comparisons test in GraphPad Prism. Results were considered statistically significant at p < 0.05. All statistics are reported as average +/- standard error of the mean, unless otherwise noted. For differential expression analysis, statistical analysis was performed entirely in R Studio with a cutoff value of p < 0.05.

## RESULTS

### Macrophage activation directly affects cell morphology

RAW 264.7 macrophages were stimulated to either an M1 phenotype (100ng/mL LPS and 20ng/mL IFN-γ) or and M2 phenotype (30ng/mL IL-4) for 24 hours. Phenotypes were confirmed using M1 (CD86 and iNOS) and M2 (CD206 and CD163) markers (Fig. 1a-e). Cell cytoskeleton staining was used to assess cell morphology. Cells cultured in polarization media showed higher area, perimeter, and length than unstimulated controls. Additionally, activated cells had lower circularity (p < 0.0001) and higher elongation (p = 0.0024) than controls. M1 polarized macrophages tended to have higher cell area (p < 0.0001) and perimeter (p < 0.0001) than both controls and M2 macrophages, while M2 polarized macrophages took on a more elongated shape (p < 0.0001) than the other groups (Fig. 1f-j).

**Figure 1.**
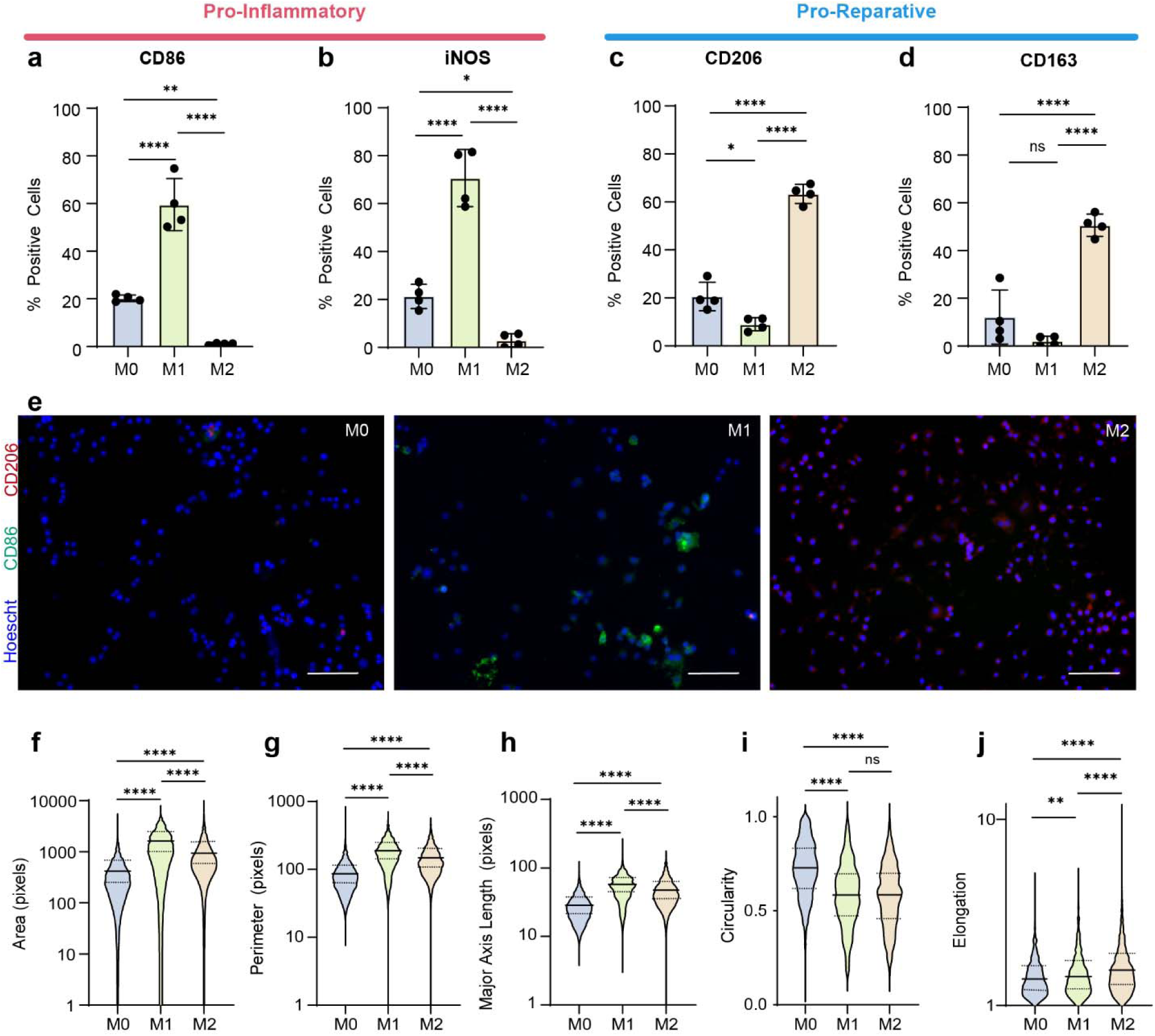
Classically and alternatively activated macrophages show distinct changes to cell morphology. Percent of macrophages expressing (a) CD86, (d) iNOS, (c) CD206, and CD163 (n = 4) when unstimulated (M0), classically activated (M1), or alternatively activated (M2), as determined by fluorescent imaging (e). Cytoskeletal staining allowed for measurement of cell (f) area, (g) perimeter, (h) major axis length, (i) circularity, and (j) elongation. Graphs display mean +/- standard error. * p < 0.05, ** p < 0.01, *** p < 0.001, **** p < 0.0001. ns denotes no significance. Scale bars = 100μm.

### ECM promotes macrophage attachment and activation

RAW 264.7 macrophages were seeded onto cECM-coated surfaces or uncoated controls and cultured for 24 hours. Neonatal cECM promoted cell attachment to the surface (p = 0.0003) and cell proliferation (p = 0.0008) compared to other groups (Fig. 2). Additionally, cells grown on all cECM coated surface showed higher levels of activation than control groups, as shown by increased cell area, perimeter, and length, and decreased cell circularity (p < 0.0001). The enhanced attachment and proliferation observed on neonatal ECM suggest that developmental age-dependent matrix composition provides a more permissive microenvironment for macrophage survival and expansion. These findings are consistent with previous reports demonstrating that tissue-derived ECM can regulate macrophage behavior through both biochemical and biophysical signaling cues (^22^, (Sadtler, 2016)).

**Figure 2.**
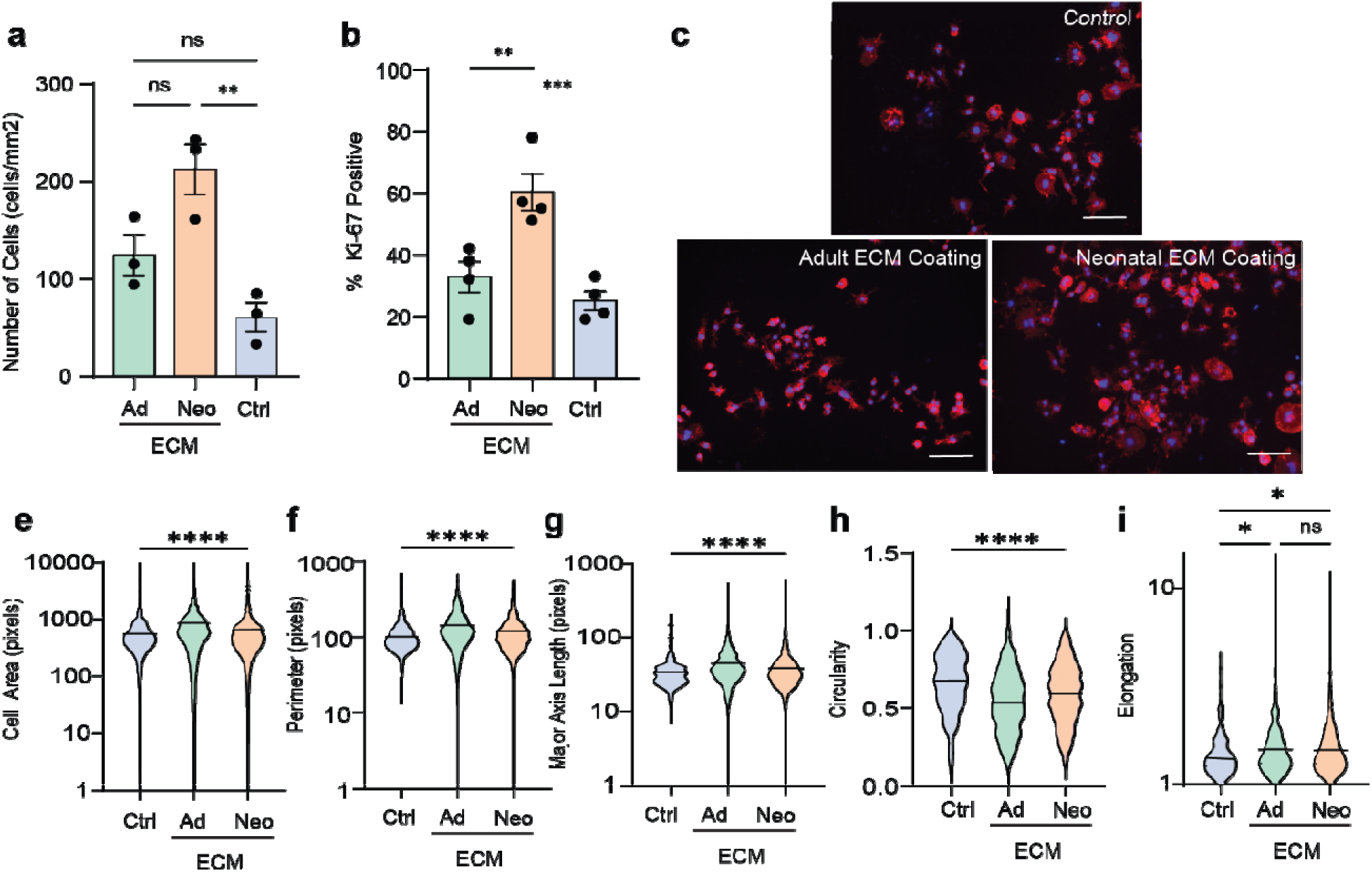
ECM coatings promote cell attachment and proliferation. (a) Cell Density (n = 4) and (b) percent of cells expressing the mitosis marker Ki-67 (n = 4). (c) Cytoskeletal staining allowed for the measurement of cell (d) area, (e) perimeter, (f) major axis length, (g) circularity, and (h) elongation. Graphs display mean +/- standard error. ** p < 0.01, *** p < 0.001, **** p < 0.0001. ns denotes no significance. Scale bars = 100μm.

### Younger developmental age ECM promotes an M2-like phenotype

Macrophage polarization after 24 hours was measured by the expression of M1 markers CD86 and iNOS and M2 markers CD206 and CD163. Expression of inflammatory M1 markers were relatively low across all groups, while expression of pro-reparative M2 markers increased with younger developmental age ECM coatings (Fig. 3). Macrophages grown on neonatal ECM coatings exhibited significantly increased expression of CD206 compared to adult ECM and control groups (p = 0.0239 and p < 0.0001), and significantly increased CD163 expression (p < 0.01) compared to all other other groups, suggesting a reparative phenotype shift. CD163 expression displayed a gradual increase in expression with younger ECM age groups. Importantly, macrophages grown on neonatal ECM showed similar levels of expression of CD206 and CD163 as macrophages stimulated to M2-like polarization with IL-4 (Fig. 1).

**Figure 3.**
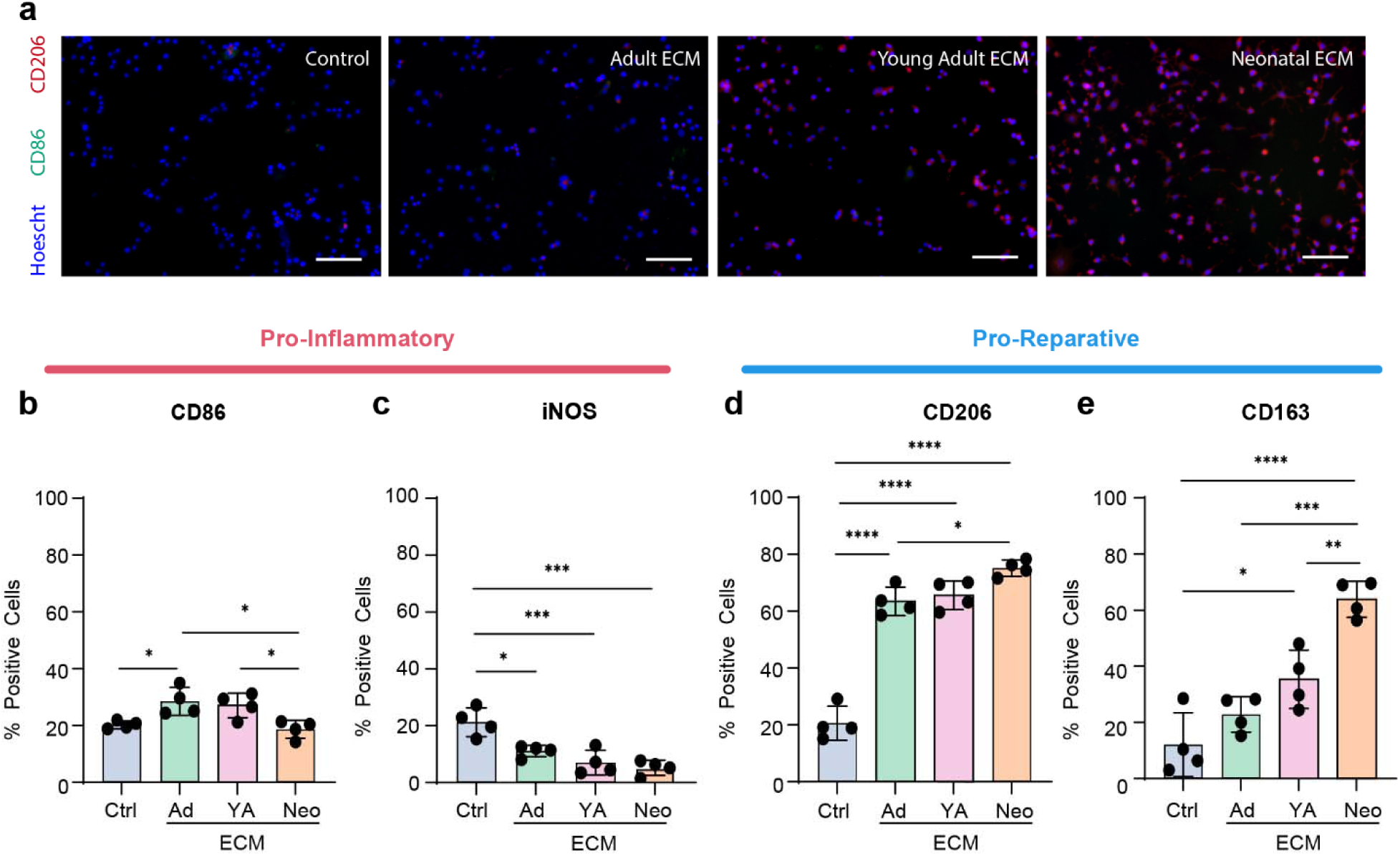
ECM promotes stronger expression of M2 markers in a developmental age-dependent manner. (a) Representative fluorescent images used to determine the percentage of cells expressing (b) CD86, (c) iNOS, (d) CD206, and (e) CD163. Graphs display mean +/- standard error. * p < 0.05, ** p < 0.01, *** p < 0.001, **** p < 0.0001. Scale bars = 100μm.

After 24 hours of culture on ECM coatings, macrophages were stimulated towards an M1 phenotype using inflammatory stimulation, as would be seen in a wound healing environment, for 24 hours and polarization was measured. Neonatal cardiac ECM decreased the expression of M1 markers compared to the M1 control, while still promoting strong M2-like expression (Fig. 4). Both CD86 and iNOS expression was significantly lower in cells grown on neonatal ECM than both adult ECM or the M1 control (p < 0.05). Additionally, CD206 and CD163 expression remained high, with CD163 expression showing the same increasing trend with decreasing ECM developmental age.

**Figure 4.**
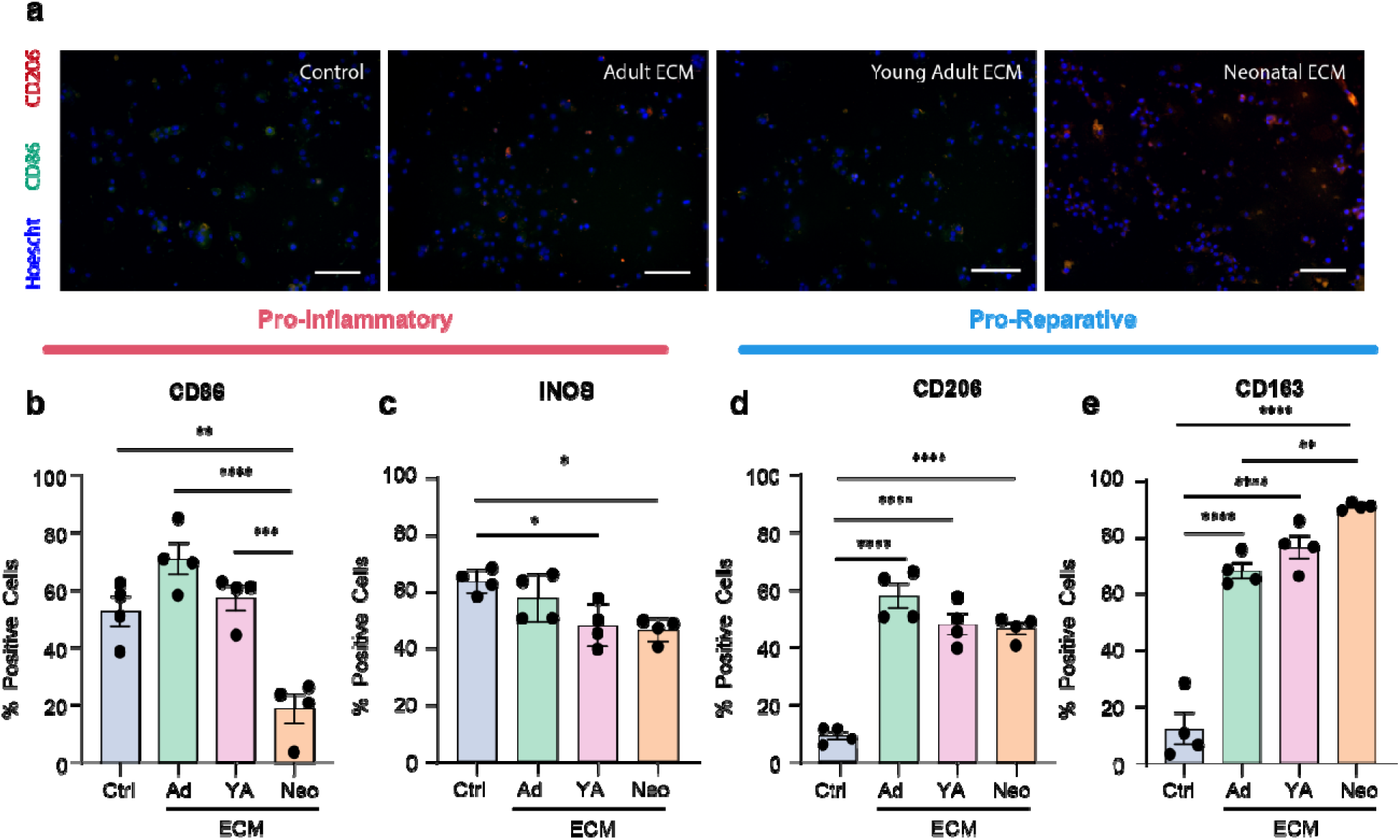
**ECM stabilizes macrophage inflammatory activation in the presence of LPS and IFN-**γ **in a developmental age-dependent manner.** (a) Representative fluorescent images used to determine the percentage of cells expressing (b) CD86, (c) iNOS, (d) CD206, and (e) CD163 (n = 4). Graphs display mean +/- standard error. * p < 0.05, ** p < 0.01, *** p < 0.001, **** p < 0.0001. Scale bars = 100μm.

### RNA Sequencing

Macrophages grown on ECM coatings or in polarization media for 24 hours underwent RNA sequencing and were compared to an unstimulated control. Compared to the M0 control, 69, 76, 10, 156, and 50 genes were differentially regulated in the M1, M2, Adult ECM, Young Adult ECM, and Neonatal ECM groups, respectively (Fig. 5a). Clustering showed stronger similarities to macrophages stimulated with either M1 or M2 polarization media than with cells cultured on ECM coatings, suggesting a unique phenotype that does not fit into either category (Fig. 5c). PCA analysis also showed highest variance in the neonatal ECM group compared to other groups. Differences in the second principal component separated out the M1 and M2 phenotypes from the ECM phenotypes. Additionally, hierarchical clustering clumped the experimental M1 and M2 phenotypes together before including any ECM phenotypes.

**Figure 5.**
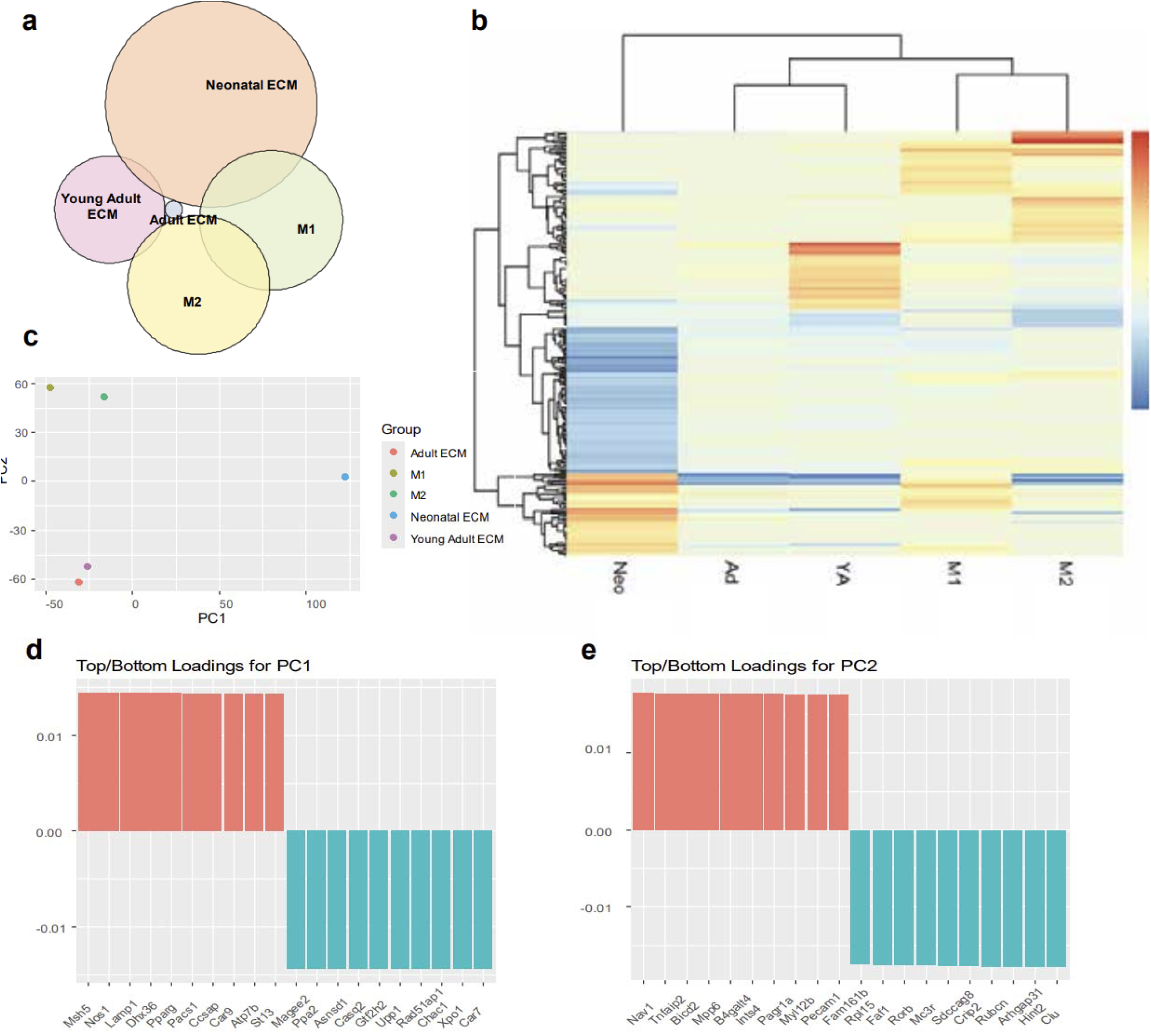
RNA sequencing reveals transcriptional differences between macrophages cultured on different ECM coatings. (a) Venn Diagram of differentially regulated genes in each group. (b) PCA analysis. (c) hierarchical heat map. (d) top and bottom loadings for the first and second principal components, as determined by PCA analysis. Significantly regulated genes determined by a fold change greater than 1 or less than -1 from M0.

Gene enrichment analysis revealed distinct differences in genes differentially regulated by different developmental age ECM (Supp. Fig. 1). Genes significantly upregulated in macrophages grown on adult ECM related largely to cell metabolism, with “ATP synthesis coupled electron transport,” “electron transport chain,” “oxidative phosphorylation,” and “proton transmembrane transport” being among the top upregulated GO terms. The young adult ECM seemed to promote translation-related gene expression, with top upregulated GO terms including “tRNA metabolic process,” “tRNA aminoacylation for protein translation,” “amino acid activation,” “protein targeting to ER,” and “establishment of protein localization to endoplasmic reticulum.” Macrophages grown on neonatal ECM prioritized genes relating to muscle development and contraction, such as “muscle organ development,” “muscle contraction,” and “actin-filament based movement.” Interestingly, these GO terms were among the most downregulated in both the adult and young adult ECM groups. Notably, a substantial proportion of the most significantly downregulated pathways in neonatal ECM-derived macrophages were associated with interferon signaling. GO terms including “response to interferon-beta”, “interferon-mediated signaling pathway”, “defense response to virus”, and “regulation of type I interferon production” demonstrated coordinated suppression, suggesting that developmental age-dependent ECM cues regulate a convergent inflammatory signaling axis.

### Neonatal ECM Downregulates Macrophage Inflammatory Response to Interferon Molecules

Because interferon signaling emerged as one of the most consistently altered biological processes identified by transcriptomic analysis, subsequent studies focused on determining whether neonatal ECM altered macrophage responsiveness to exogenous inflammatory stimulation. To assess this, macrophages were grown on ECM coatings for 24 hours before being introduced to either IFN-γ or IFN-β alone. Polarization was then assessed by measuring CD206 and CD86 expression. Macrophages stimulated by IFN-γ showed moderate levels of CD86 expression, which decreased only in the presence of neonatal ECM (p < 0.05). Macrophages cultured on ECM matrikines still maintained comparable levels of CD206 to previous data (Fig. 6). IFN-β resulted in a stronger inflammatory response, as seen at increased expression of CD86, which was again decreased only in the presence of neonatal ECM (p < 0.0001) (Fig. 7). While inflammatory stimulation shifted macrophages toward a more activated phenotype across all groups, macrophages cultured on neonatal ECM maintained expression profiles more consistent with a reparative state than those cultured on young adult, adult, or uncoated substrates. These findings suggest that neonatal ECM not only influences basal macrophage phenotype but may also confer resistance to inflammatory reprogramming.

**Figure 6.**
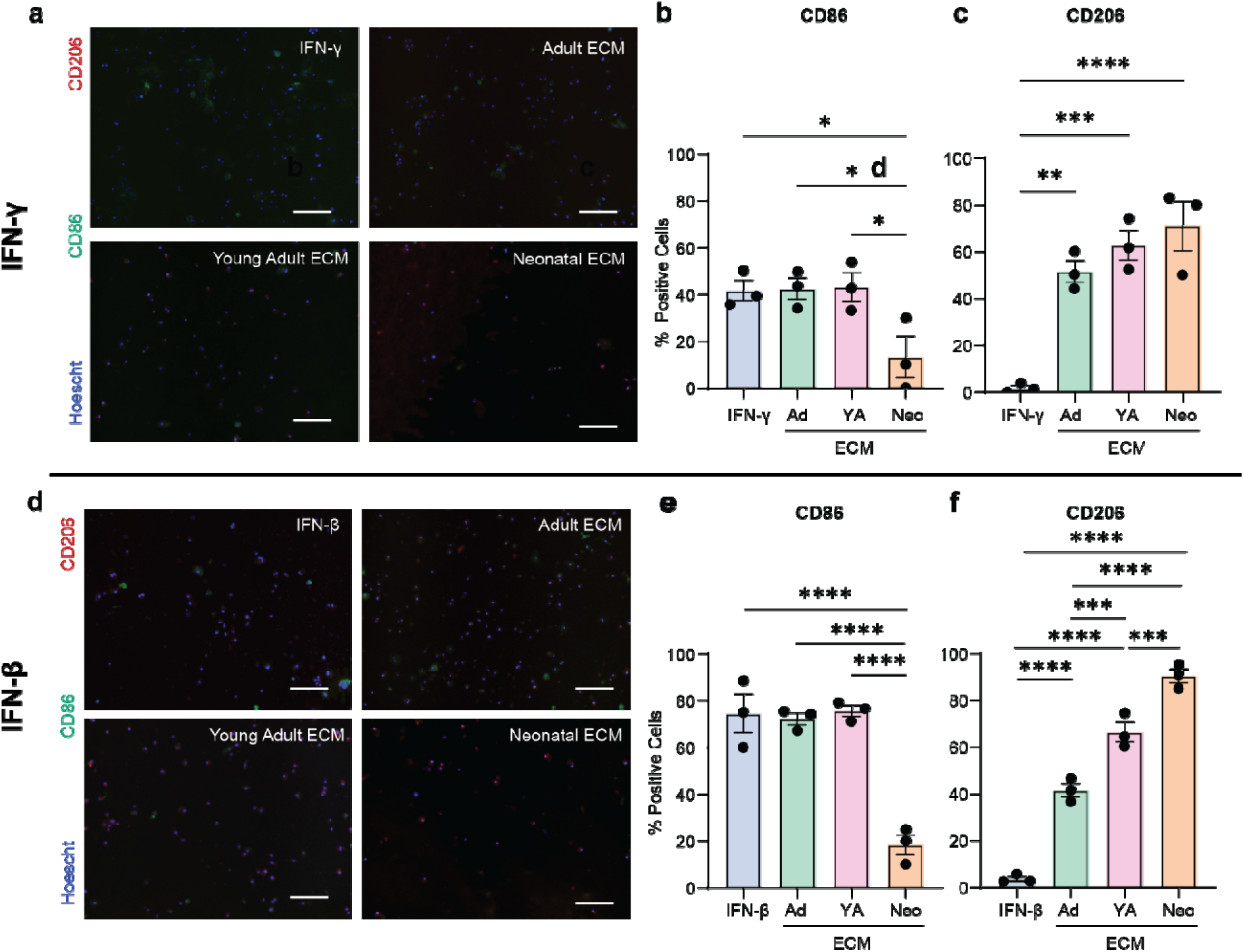
Interferon induced activation was significantly downregulated in the presence of neonatal ECM. (a) Representative fluorescent images of macrophages grown on ECM coatings in the presence of IFN-γ, as used to determine the percent of cells expressing (b) CD86 and (c) CD206 (n = 3). (d) Representative fluorescent images of macrophages grown on ECM coatings in the presence of IFN-β, as used to determine the percent of cells expressing (e) CD86 and (f) CD206 (n = 3). Graphs display mean +/- standard error. *** p < 0.001, **** 0 < 0.0001. Scale bars = 100μm.

**Figure 7.**
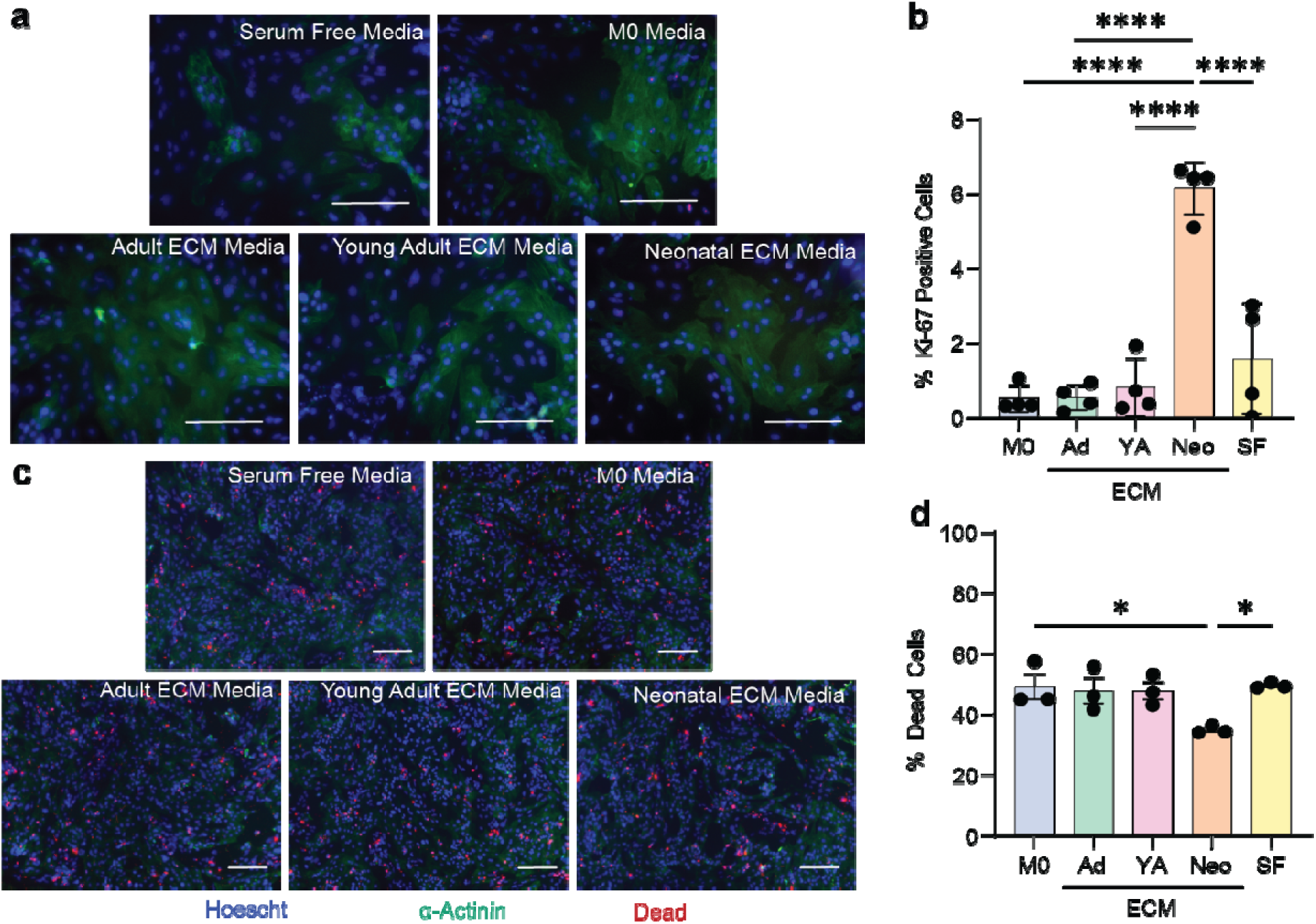
Macrophage secretions influence iPSC derived CM activity. (a) Representative images of iPSC-CM cultured with macrophage conditioned media, as used to determine (b) the percent of α-actinin positive cells expressing the mitosis marker Ki-67 (n=4). (c) Representative images of (d) iPSC-CM survival after exposure to oxidative stress (n=3). Graphs display mean +/- standard error. * p < 0.05, ** p < 0.01, **** p < 0.0001. Scale bars = 100μm.

### Macrophage Secretome can Influence Other Cardiac Cell Types

To determine if macrophage phenotype expression could influence other cells in the cardiac wound healing environment, macrophage conditioned media from cells grown on different ECM coatings, or an uncoated control, were collected and used to culture iPSC derived cardiac fibroblasts and cardiomyocytes. Only conditioned media collected from macrophages grown on neonatal cECM coatings promoted an increase in cardiomyocyte proliferation (p < 0.0001) (Fig. 8a). Additionally, media from macrophages grown on neonatal cECM coatings were able to rescue oxidatively stressed cardiomyocytes compared to other groups (p < 0.05) (Fig. 8b). While no changes in cardiac fibroblast proliferation were seen between the different ECM groups, neonatal cECM promoted cardiac fibroblast activation independent of TGF-β (p < 0.001) (Fig. 8c-d). Together, these findings indicate that developmental-age ECM influences both transcriptional and secretory aspects of macrophage activation. The coordinated attenuation of inflammatory cytokine production further supports the transcriptomic evidence for suppressed inflammatory signaling in neonatal ECM-conditioned macrophages.

**Figure 8.**
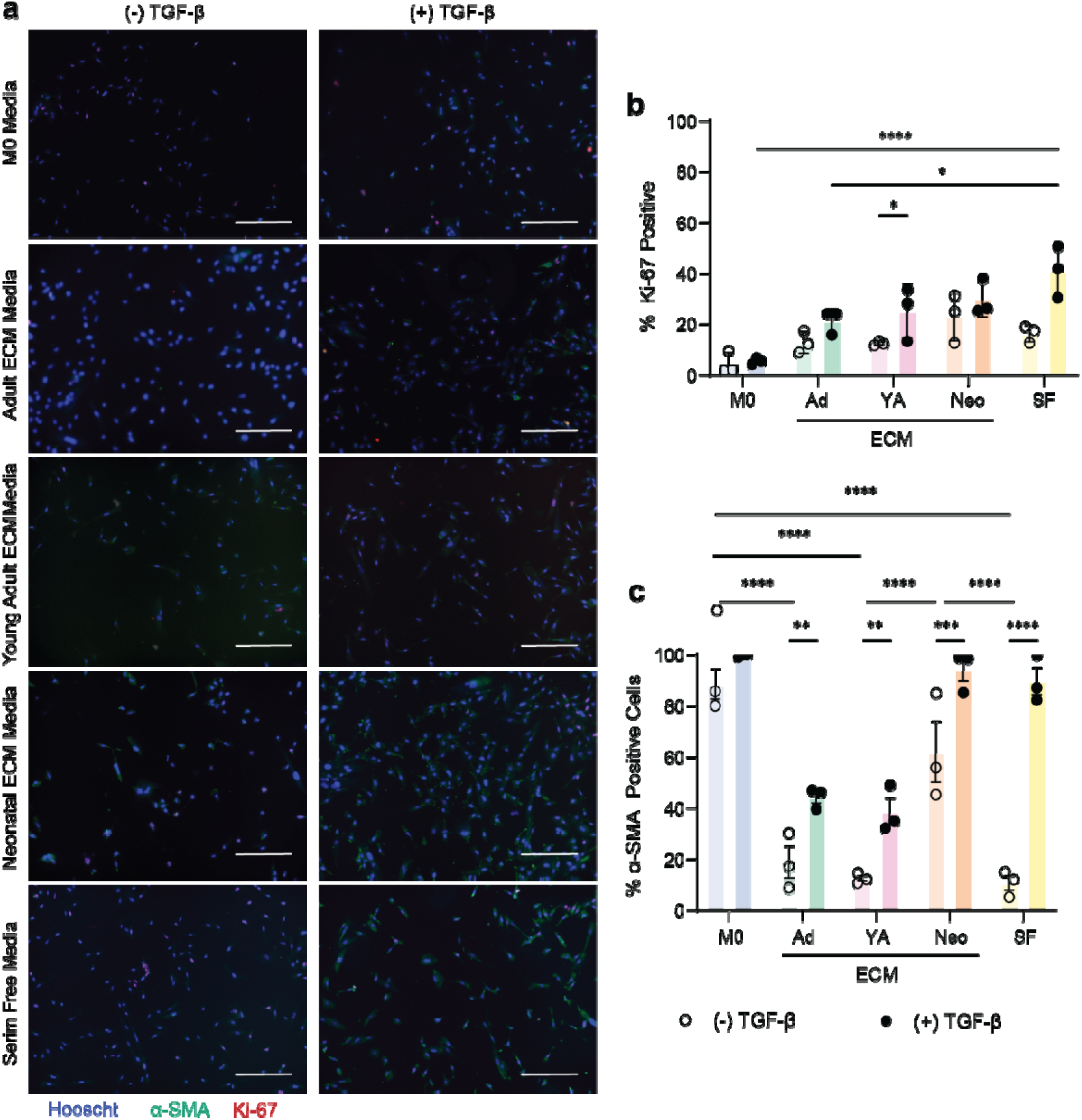
Macrophage secretions influence iPSC derived CF activity. (a) Representative images of iPSC-CF cultured using macrophage conditioned media with and without of the addition of TGF-β as used to determine the expression of (b) Ki-67 and (c) αSMA (n = 3). Graphs display mean +/- standard error. * p < 0.05, ** p < 0.01, *** p < 0.001, **** p < 0.0001. Scale bars = 100μm.

## Discussion

Tissue remodeling, during normal tissue homeostasis or wound healing, is initiated by extracellular matrix degradation, where ECM proteins are broken down into small peptides that have the ability to give unique signals to cells relating to tissue remodeling. There is increasing evidence in the literature that signals derived from these peptides can influence cell behavior ^12^, including macrophage phenotype ^13^. As ECM composition changes with organism developmental age ^12^, so do the signals generated. An abundance of evidence from the literature suggests that source animal age greatly impacts how extracellular matrix stimulates cells (^11, 12, 13, 23^), with ECM from neonatal and fetal animals generally having a highly beneficial effect on cells ^12^. This study demonstrates that the developmental age of cardiac extracellular matrix substantially alters macrophage phenotype and inflammatory responsiveness. While neonatal ECM promoted expression of reparative macrophage markers under basal conditions (Fig. 3), transcriptomic analysis revealed that these cells do not represent a classically polarized M2 population (Fig. 5). Instead, macrophages exposed to neonatal ECM exhibited a distinct activation state characterized by suppression of interferon-associated inflammatory pathways and reduced responsiveness to subsequent inflammatory stimulation (Fig. 6 and 7). In addition, these findings support the emerging concept that extracellular matrix functions as a reservoir of developmental memory. Although the original matrix-producing cells are absent following decellularization, age-specific biological information remains encoded within the ECM and is sufficient to alter macrophage behavior. Such developmental memory has previously been reported for cardiac ECM effects on fibroblasts ^24^, stem cells ^25^, and cardiomyocytes ^12^; the current work extends this paradigm to innate immune regulation.

The increase in pro-regenerative protein expression in macrophages cultured on ECM coated substrates has been previously documented (^22, 26^). This M2 shift makes sense, since matrikines are naturally released during the maintenance of tissue homeostasis, a process which is well known to be supported by M2-like tissue resident macrophages ^27^. The increase in M2 expression in macrophages grown on neonatal ECM suggests that microenvironmental signaling from tissue remodeling may play a role in the regenerative capacity of neonatal tissue *in vivo*. Likewise, the increase in proliferation of macrophages culture on neonatal ECM may play a role in the local proliferation of tissue resident macrophages following injury in neonatal animals not found in adult tissues.

Macrophages cultured on neonatal ECM were also able to maintain this M2 expression in the presence of inflammatory stimulation, as seen with high CD206 and CD163 expression, while showing decreased expression of M1 markers (Fig. 4). The ability of younger ECM to attenuate iNOS expression has been previously documented ^13^, but not among the more regenerative neonatal ECM. The drop in iNOS signaling between 6 week old ECM, which is typically seen in the literature, and neonatal ECM, shows the importance of utilizing signals from highly regenerative neonatal ECM.

RNA sequencing revealed a significant downregulation of several players in the interferon signaling pathway (Fig. 5, Supp. Fig. 1). While interferon signaling was originally characterized in antiviral immunity, accumulating evidence demonstrates important roles for interferon-dependent pathways in sterile tissue injury and cardiac remodeling. Following myocardial infarction, release of intracellular nucleic acids activates innate immune signaling cascades that converge on IRF-dependent transcriptional programs and type I interferon production. Excessive activation of these pathways contributes to persistent inflammatory signaling and adverse remodeling (^28, 29, 30^). Confirming this important, role for Interferon signaling in post-MI remodeling, recent studies have found that inhibiting type I interferon signaling decreased the presence of inflammatory cytokines, reduced monocyte infiltration, and improved long term cardiac function in mice ^28^. Additionally, neonates have previously been shown to be ineffective at initiating inflammation via type I interferon signaling, a characteristic that has previously been exclusively associated with immature cells ^31^. This study suggests that microenvironmental signaling may play a role in ineffective immune responses in neonatal mammals, and that suppression of interferon-associated pathways observed in neonatal ECM-conditioned macrophages may represent a therapeutically relevant mechanism by which regenerative cardiac matrices attenuate maladaptive inflammation.

This also suggests that the use of LPS to stimulate macrophages towards an inflammatory phenotype may confound results when studying sterile injuries. LPS is well known to cause a robust and well characterized M1 reaction in macrophages, utilizing specific pathways for bacterial infections. However, inflammatory stimulation during sterile injury utilizes a different subset of signaling pathways, which may be up or down regulated differently. Future studies will utilize more relevant inflammatory stimuli, such as extracellular oligonucleotides.

While this study investigates changes in macrophage phenotype and activation, changes to macrophage function were not directly investigated. Instead, the ability of macrophage secretions to influence other cells common in the cardiac wound healing environment was studied. Both cardiomyocyte and cardiac fibroblast proliferation was seen to increase in the presence of secretions from macrophages grown on neonatal ECM (Fig. 8), suggesting an increase in growth factor release. These results suggest that macrophages may stimulate cardiomyocyte recovery and matrix remodeling in the neonatal cardiac wound healing environment. This conclusion is supported by studies that show macrophages are necessary for proper wound healing in neonates, and that removal of macrophages leads to fibrosis ^9^. Contrary to the necessity of macrophages for neonatal wound healing, other studies have shown that monocyte derived macrophage recruitment may be detrimental to healing in adult cardiac wound healing ^32^. This suggests an origin-dependent response in macrophages to different signaling environments.

Several limitations should be acknowledged. As previously noted, the current study used RAW264.7 macrophages, which may not completely recapitulate primary macrophage responses. Indeed, previous studies have shown that THP-1 macrophages, an immortalized monocyte cell line, and primary bone marrow derived macrophages respond different to ECM stimulation ^22^, highlighting the importance of continued investigation of macrophage source in this work. In addition, .in our cytokine secretion assay we used iPSC derived cardiac fibroblasts and cardiomyocytes, which can be considered immature.. Lastly, while transcriptomic analyses identified interferon signaling as a major regulated pathway, additional mechanistic studies will be necessary to identify the specific ECM components responsible for this effect. Future studies using primary macrophages, adult cells in vitro, in vivo studies, proteomic characterization of ECM composition, and direct modulation of interferon signaling pathways will further clarify these mechanisms.

In summary, in this study, we investigated how the developmental age of cardiac extracellular matrix influences macrophage phenotype and function. Macrophages cultured on neonatal cardiac ECM exhibited enhanced attachment and proliferation, increased expression of reparative macrophage markers, and altered transcriptional profiles compared to cells cultured on young adult ECM, adult ECM, or uncoated substrates. Transcriptomic analysis further revealed significant developmental age-dependent differences in inflammatory pathway regulation, particularly within interferon-associated signaling networks, which were accompanied by attenuated responses to subsequent inflammatory stimulation. Collectively, these findings demonstrate that developmental age-dependent cardiac ECM regulates macrophage phenotype and inflammatory responsiveness. Neonatal cardiac ECM promotes a distinct reparative activation state characterized by suppression of interferon-associated inflammatory signaling, identifying a potential immunomodulatory mechanism contributing to the pro-regenerative properties of neonatal cardiac extracellular matrix. These findings further support the concept that extracellular matrix serves as a reservoir of developmental information capable of directing immune cell behavior and suggest that age-specific ECM cues may represent an important strategy for engineering immunomodulatory biomaterials for cardiac repair and regeneration.

